# High-quality genome assembly and high-density genetic map of asparagus bean

**DOI:** 10.1101/521179

**Authors:** Qiuju Xia, Ru Zhang, Xuemei Ni, Lei Pan, Yangzi Wang, Xiao Dong, Yun Gao, Zhe Zhang, Ling Kui, Yong Li, Wen Wang, Huanming Yang, Chanyou Chen, Jianhua Miao, Wei Chen, Yang Dong

## Abstract

Asparagus bean (*Vigna*. *unguiculata* ssp. *sesquipedialis*), known for its very long and tender green pods, is an important vegetable crop broadly grown in the developing countries. Despite its agricultural and economic values, asparagus bean does not have a high-quality genome assembly for breeding novel agronomic traits. In this study, we reported a high-quality 632.8 Mb assembly of asparagus bean based on the whole genome shotgun sequencing strategy. We also generated a high-density linkage map for asparagus bean, which helped anchor 94.42% of the scaffolds into 11 pseudo-chromosomes. A total of 42,609 protein-coding genes and 3,579 non-protein-coding genes were predicted from the assembly. Taken together, these genomic resources of asparagus bean will facilitate the investigation of economically valuable traits in a variety of legume species, so that the cultivation of these plants would help combat the protein and energy malnutrition in the developing world.

## Background & Summary

Asparagus bean (*Vigna unguiculata* ssp. *sesquipedialis*, 2n = 2 x =22) is a warm-season and drought-tolerant subspecies of cowpea (*Vigna unguiculata*) with a wide cultivation area in East and Southeast Asia^1^. This plant is also known as yardlong bean because of its characteristic pod that grows up to 50-100 cm in length^2^. The long pod trait is believed to be the result of intensive local domestication after it was brought to Asia from sub-Saharan Africa^3^. Unlike the grain-type subspecies common cowpea (*Vigna*. *unguiculata* ssp. *unguiculata*, or black-eyed pea), asparagus bean is harvested while its pod is still tender, thereby providing a very good source of protein, minerals, vitamins, and dietary fiber^4^. Due to the low requirement for cultivation management and its high nutritional value, asparagus bean is one of the top crops that help combat malnutrition and food insecurity in most developing countries^5^.

As the DNA sequencing technologies became more advanced and affordable for the past decade, previous research had mainly focused on delineating the genome of common cowpea (estimated genome size of 620 Mb^6^). The first common cowpea (variety IT97K-499-35) genomic resources included a partial 323 Mb whole-genome shotgun assembly^7^, a 497 Mb bacterial artificial chromosome physical map^7^, and consensus genetic maps based on either 10K^8^ or 50K single nucleotide polymorphisms (SNPs)^7^. A more recent research reported two survey genomes of common cowpea (varieties IT97K-499-35 and IT86D-1010) with substantially improved assembly sizes (568 Mb and 609 Mb, respectively)^9^. In addition, a draft IT97K-499-35 variety reference genome was assembled by incorporating the single molecule real-time technology, yielding an assembly size of 519.4 Mb into 722 scaffolds and 11 pseudo-chromosomes (http://phytozome.jgi.doe.gov/). In comparison, the genetic resources for asparagus bean are lacking despite its agricultural and economic importance. So far, only three genetics maps were derived from either simple sequence repeat markers^10,11^ or restriction-site associated DNA sequencing for asparagus bean^12^.

In this study, we aimed to fill the knowledge gap with regard to the asparagus bean genome and provide new genetic resources for breeding cowpea and related legume species. A schematic workflow of the research was showcased in Fig. 1. In brief, a series of short-insert and large-insert Illumina libraries were sequenced on an Illumina HiSeq 4000 platform, yielding a total of 222.9 Gb clean data (Table 1). Since the genome size of asparagus bean was estimated to be 652.4 Mb using the *K*-mer distribution analysis (Fig. 2), the clean data used for genome assembly represented about 340 x coverage. The software SOAPdenovo^13^ was used to generate a draft contig assembly of 549.8 Mb with a contig N50 size of 15.2 kb (Table 2). After scaffolding and gap closing, the final asparagus genome was 632.8 Mb (96.98% of the estimated genome) in size with scaffold N50 size of 2.7 Mb (Table 2, Data Citation 1). We also obtained 536,824 high-confident SNPs from the whole-genome sequencing data of 97 asparagus bean F2 individuals and two parents from a well-controlled selfing population. These SNPs were used to construct a high-density genetic map for asparagus bean, in which 1,556 scaffolds were successfully anchored onto 11 pseudo-chromosomes (Table 3). Furthermore, the asparagus bean genome contained 294.95 Mb of transposable elements, accounting for 46.47% of the assembly (Table 4 and 5). The gene prediction was performed on a combination of *de novo*, homologous, and RNA-Seq-based approaches. It resulted in 42,609 protein-coding genes and 3,579 non-protein-coding genes, respectively (Table 6).

**Figure 1.**
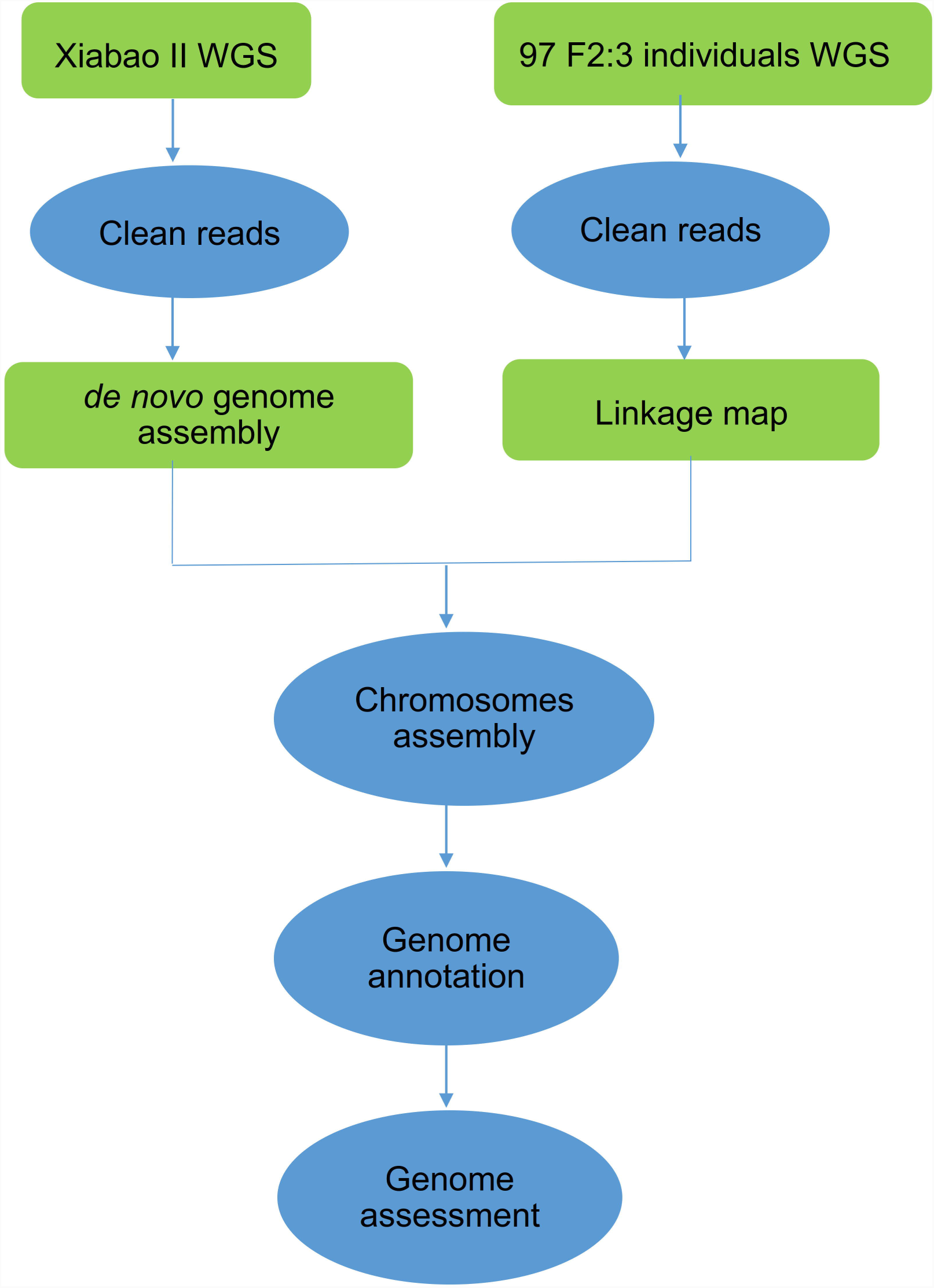
General description of the assembly workflow. The pipeline included removal of low quality and adapter-contaminated reads, *de novo* assembly, construction of linkage map, chromosome-scale assembly, and genome annotation.

**Figure 2.**
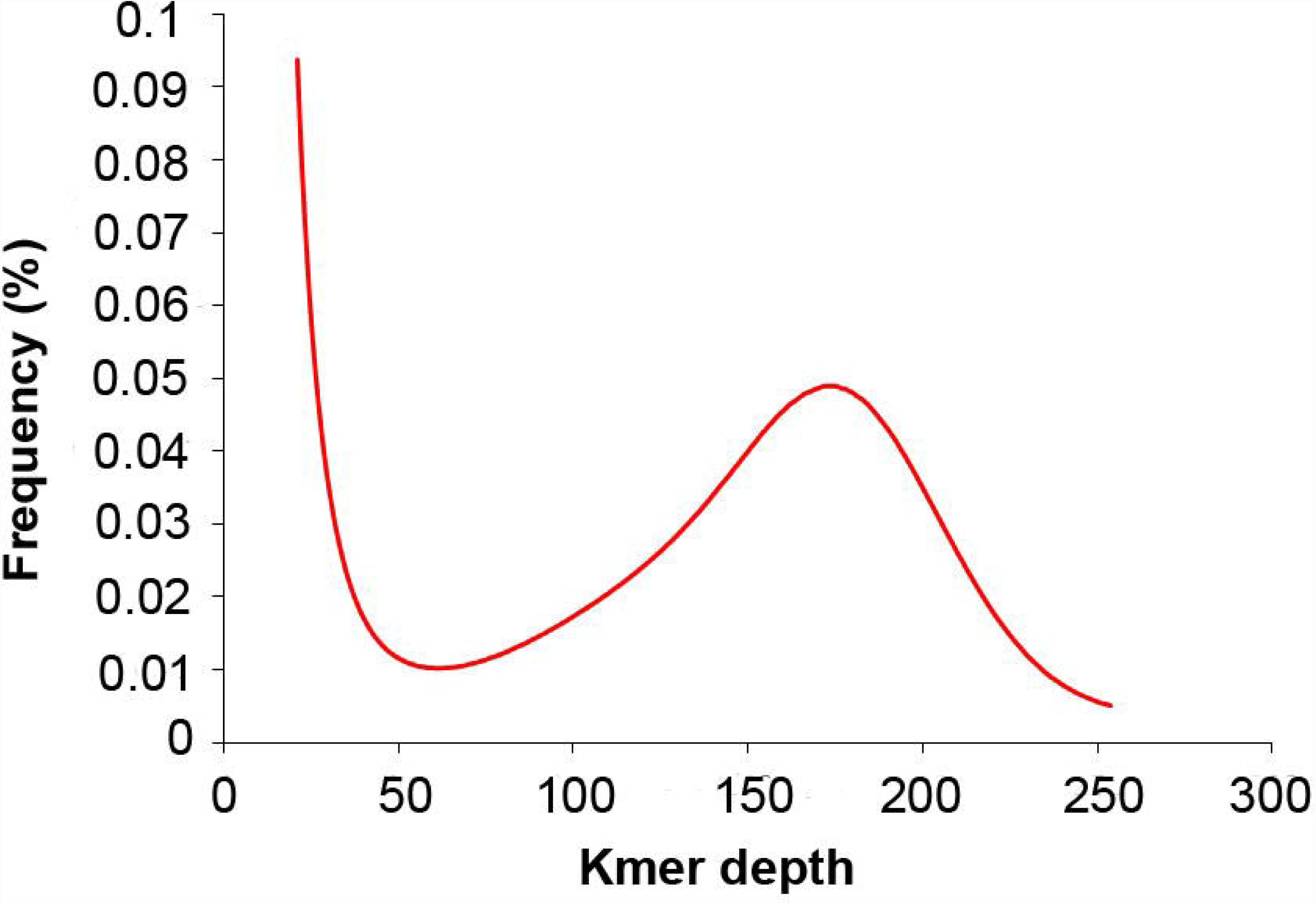
17-mer frequency distribution of sequencing reads.

**Table 1.** Statistics of raw data after filtering.

| Insert Size | Clean Length (bp) | Number of Clean Reads | Clean Bases (Gb) | Sequence Coverae (X) |
| --- | --- | --- | --- | --- |
| 350 | 2 X 125 #Hiseq4000 | 143,324,095 | 35.831 | 54.92 |
| 445 | 2 X 125 #Hiseq4000 | 200,584,850 | 50.146 | 76.86 |
| 758 | 2 X 125 #Hiseq4000 | 60,211,855 | 15.053 | 23.07 |
| 912 | 2 X 125 #Hiseq4000 | 113,659,706 | 28.415 | 43.55 |
| 2000 | 2 X 125 #Hiseq4000 | 79,141,602 | 19.785 | 30.32 |
| 3000 | 2 X 125 #Hiseq4000 | 82,610,562 | 20.653 | 31.65 |
| 5000 | 2 X 125 #Hiseq4000 | 80,415,362 | 20.104 | 30.81 |
| 9000 | 2 X 125 #Hiseq4000 | 72,037,228 | 18.009 | 27.6 |
| 15000 | 2 X 125 #Hiseq4000 | 59,701,495 | 14.925 | 22.87 |
| Total | - | 891,686,755 | 222.921 | 341.66 |

**Table 2.**
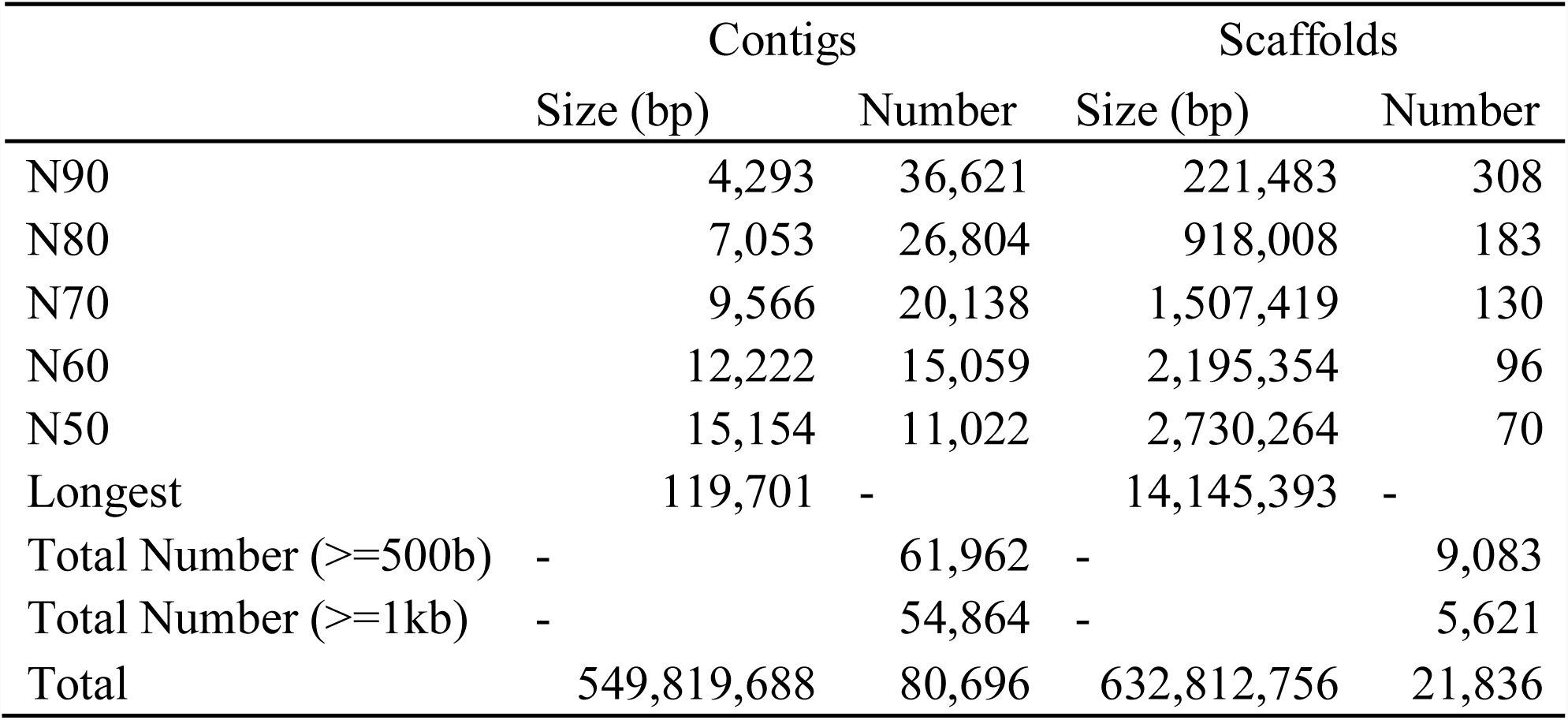
Results of the asparagus bean genome assembly.

|  | Contigs |  | Scaffolds |  |
| --- | --- | --- | --- | --- |
|  | Size (bp) | Number | Size (bp) | Number |
| N90 | 4,293 | 36,621 | 221,483 | 308 |
| N80 | 7,053 | 26,804 | 918,008 | 183 |
| N70 | 9,566 | 20,138 | 1,507,419 | 130 |
| N60 | 12,222 | 15,059 | 2,195,354 | 96 |
| N50 | 15,154 | 11,022 | 2,730,264 | 70 |
| Longest | 119,701 | - | 14,145,393 | - |
| Total Number ($\geq 500$b) | - | 61,962 | - | 9,083 |
| Total Number ($\geq 1$kb) | - | 54,864 | - | 5,621 |
| Total | 549,819,688 | 80,696 | 632,812,756 | 21,836 |

**Table 3.** Statistics of pseudo-chromosomes and genetic map in asparagus bean.

| Chromosomes | Anchored Scaffolds Number | Total length (Mb) | SNP Number | bin marker Number | Genetic distance (cM) |
| --- | --- | --- | --- | --- | --- |
| Vu1 | 97 | 45.38 | 36,916 | 154 | 94.72 |
| Vu2 | 161 | 82.25 | 58,426 | 306 | 398.24 |
| Vu3 | 224 | 54.99 | 80,711 | 162 | 214.19 |
| Vu4 | 81 | 41.88 | 41,888 | 170 | 125.31 |
| Vu5 | 233 | 55.8 | 40,719 | 175 | 185.51 |
| Vu6 | 162 | 52.07 | 54,989 | 159 | 113.72 |
| Vu7 | 148 | 49.22 | 44,186 | 193 | 333.04 |
| Vu8 | 203 | 49.61 | 95,735 | 155 | 164.48 |
| Vu9 | 81 | 51.81 | 23,748 | 189 | 207.05 |
| Vu10 | 87 | 60.58 | 31,849 | 171 | 83.74 |
| Vu11 | 79 | 53.94 | 27,657 | 179 | 260.14 |
| Total | 1556 | 597.53 | 536,824 | 2013 | 2180.14 |

**Table 4.** Statistics of repeats in the asparagus bean genome.

| Type | Repeat Size ( bp ) | % of genome |
| --- | --- | --- |
| Trf | 67,718,076 | 10.67 |
| Repeatmasker | 41,222,404 | 6.49 |
| Proteinmask | 64,741,265 | 10.2 |
| *De novo* | 264,487,557 | 41.67 |
| Total | 294,953,638 | 46.47 |

**Table 5.** TEs content in the assembled asparagus bean genome.

| Type | Rebase TEs |  | TE proteins |  | De novo |  | Combined TEs |  |
| --- | --- | --- | --- | --- | --- | --- | --- | --- |
|  | Length (bp) | % in genome | Length (bp) | % in genome | Length (bp) | % in genome | Length (bp) | % in genome |
| DNA | 6,870,914 | 1.0825 | 9,850,112 | 1.5518 | 41,887,195 | 6.5992 | 46,098,143 | 7.2626 |
| LINE | 698,393 | 0.11 | 1,195,466 | 0.1883 | 1,651,447 | 0.2601 | 2,666,968 | 0.4201 |
| SINE | 30,804 | 0.0048 | - | - | 62,452 | 0.0098 | 74,704 | 0.0117 |
| LTR | 33,989,514 | 5.3549 | 53,894,224 | 8.4908 | 112,113,184 | 17.6631 | 122,145,625 | 19.2437 |
| Other | 13,118 | 0.002 | - | - | - | - | 13,118 | 0.002 |
| Unknown | - | - | - | - | 119,021,287 | 18.7515 | 119,021,287 | 18.7515 |
| Total | 41,222,404 | 6.494 | 64,741,265 | 10.1998 | 261,567,318 | 41.2092 | 272,160,906 | 42.8782 |

**Table 6.**
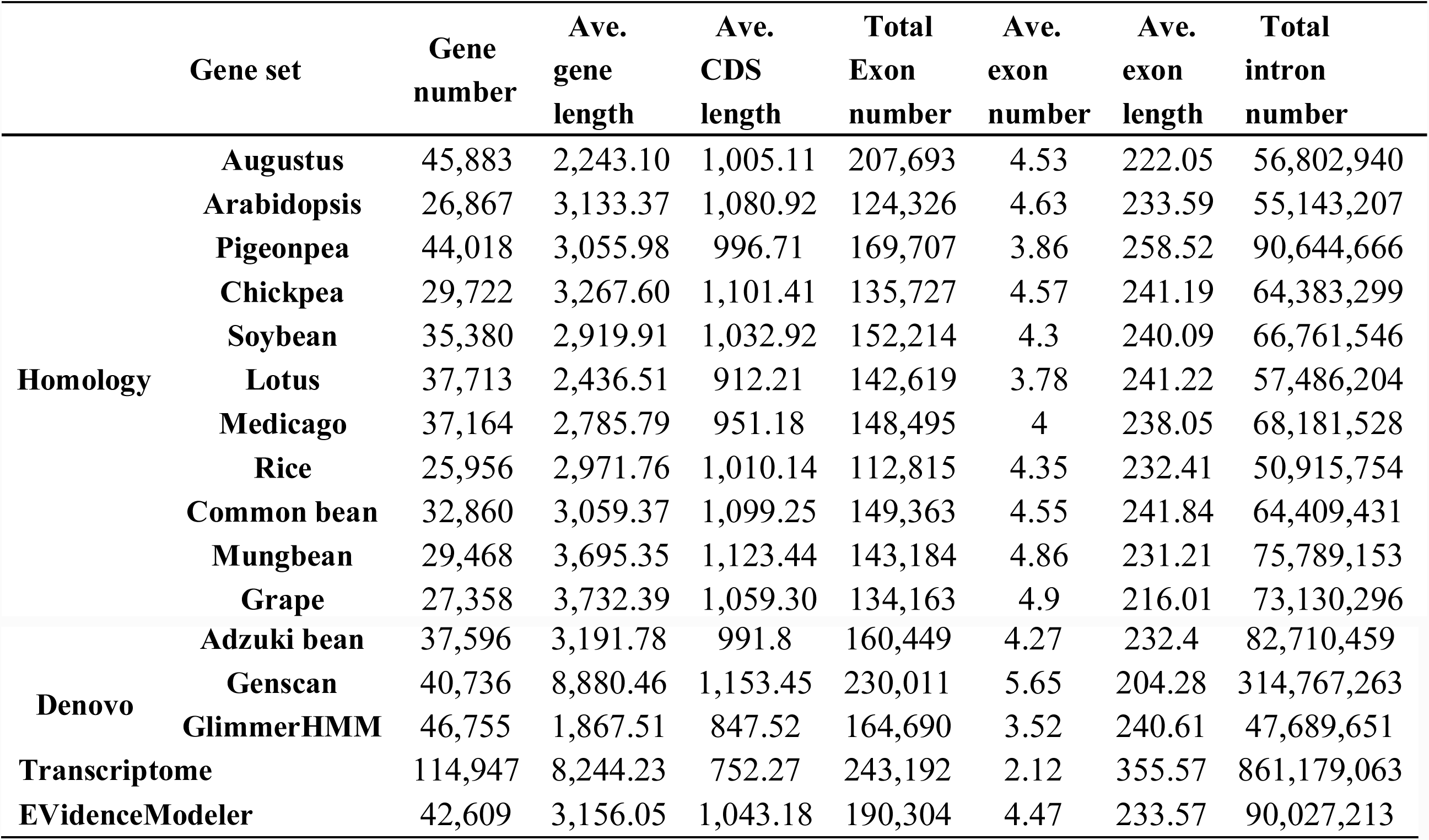
Prediction of protein-coding genes in asparagus bean genome.

## Methods

### Materials

All plant accessions were provided by Hubei Natural Science Resource Center for Edible Legumes in Wuhan of China. A single plant of the widely cultivated asparagus bean variety ‘Xiabao II’ (*Vigna unguiculata* ssp. *sesquipedialis* var. ‘Xiabao II’) was used for *de novo* sequencing and genome assembly. A F2 sequencing population was obtained for making the genetic map according to the following procedure. First, the F1 population were obtained by crossing ‘Xiabao II’ (male, same plant used for *de novo sequencing*) with a cultivar from the other subspecies, ‘Duanjiangdou’ (*Vigna unguiculata* ssp. *unguiculata* var. ‘Duanjiangdou’; female). This step yielded 17 seeds, from which only 12 seeds survived till flowering. These F1 individuals were bagged to promote selfing, which produced 561 seeds in total (the F2 generation). Only 367 of the F2 individuals were able to germinate and mature into full plants. We selected 97 of the 367 F2 individuals for genome sequencing and genetic map construction.

### Whole-genome shotgun sequencing

Young leaves were collected from a single ‘Xiabao II’ plant and used for genomic DNA extraction by the CTAB method^14^. About 10 µg of genomic DNA were used for library construction. Four short-insert libraries (350 bp, 445 bp, 758 bp, and 912 bp) and five large-insert libraries (2 kb, 3 kb, 5 kb, 9 kb, and 15 kb) were constructed with NEBNext Ultra II DNA Kit (NEB, America) and Nextera Mate Pair Sample Preparation Kit (Illumina, America), respectively. These libraries were sequenced on an Illumina HiSeq4000 platform. To ensure high-quality for the subsequent *de novo* assembly step, we filtered out the low-quality data by the following criteria: (a) reads with >2% unidentified nucleotides (N) or with poly-A structure; (b) reads with ≥40% bases having low quality for short insert-size libraries and ≥ 60% for large insert-size libraries; (c) reads with adapters or PCR duplication; (d) reads with 20 bp in 5’ terminal and 5 bp in 3’ terminal. Subsequently, about 222.9 Gb clean data were retrieved, covering 341.66-fold of the estimated genome (Table 1, Data Citation 2).

The genomic DNA was extracted with the same procedure for the parents and all 97 F2 individuals in the resequencing population. Each DNA was used to construct 500 bp insert size libraries, which were then sequenced on an Illumina HiSeq4000 platform. Each individual was sequenced to at least 4 x coverage. NGSQCToolkit_v2.3.3^15^ was used to filter low-quality reads (parameters : -l 70 -s 25) and trim the poor-quality terminal bases (parameters: -l 5 -r 5). A total of 88.26 Gb clean bases were kept, which represented 99% of the raw sequencing data (Data Citation 3).

### Estimation of the genome size

The genome size of asparagus bean was estimated by the distribution of 17-mer depth using the filtered short-insert (<1 kb) sequencing data. The peak depth of the 17-mer distribution curve was 173, and the total *K-*mer count was 112,876,121,127. The genome size was estimated to be 652.5Mb using the formula Genome_Size = Kmer_num/Peak_depth (Fig. 2).

### *De novo* genome assembly

Clean data from short insert-size libraries were corrected with the Error Correction program in SOAPdenovo package^13^. Genome assembly was performed based on the *de Bruijn* graph algorithm using SOAPdenovo package^16^ by the following steps: (1) the paired-end reads of all libraries were used to construct the contig sequences while the *K*-mer values were set as 95 and 85 at the pregraph step and map step, respectively; (2) mapped paired reads were used to construct scaffolds; (3) The GapCloser package was used to map reads to the flanking sequences of gaps and to close gaps between the scaffolds; (4) genome sequence was randomly broken to re-scaffold with SSPACE package. Gaps were then filled again by GapCloser to obtain the final assembly. In the end, there were 54,864 out of 80,696 contigs with sizes longer than 1 kb. The total length of the contig assembly was 549.81 Mb (Table 2). The longest scaffold was 14,145,393 bp, and a total of 5,621 scaffolds were longer than 1,000 bp. The total length of the scaffold assembly was 632.8 Mb (Table 2).

### High-density genetic map construction and genome assembly anchoring

All clean data obtained from the two parents and the 97 F2 individuals were mapped to asparagus bean scaffold assembly using Burrows-Wheeler-Alignment tool (BWA)^17^ mem algorithm. The SAM files were converted to BAM files using SAMtools^18^. Then the bam files were used to call SNP by the GATK software package^15^ with parameters “-T HaplotypeCaller -stand_call_conf 30.0 -stand_emit_conf 10.0” and “-T SelectVariants -selectType SNP”. The SNPs were filtered using GATK with parameters as the following: --filterExpression “QD < 2.0 ‖ ReadPosRankSum < −8.0 ‖ FS > 60.0 ‖ MQ < 40.0 ‖ SOR > 3.0 ‖ MQRankSum < −10.0 ‖ QUAL < 30” --logging_level ERROR --missingValuesInExpressionsShouldEvaluateAsFailing. After genotyping, the raw SNPs were filtered with the following criteria: missing rate <0.3 and heterozygous genotypes <0.5, resulting in a total of 836,933 high-confidence SNPs.

For the genetic map construction, 50 SNPs were selected to generate bin markers from the two termini and middle part of each scaffold. These bin markers were grouped into 11 linkage groups by JoinMap v4.1^19^ with the regression mapping algorithm. The grouped bins were then sorted and genetic distance was calculated by MSTmap with the Kosambi model^20^. According to this linkage map, scaffolds were anchored onto 11 pseudo-chromosomes. The SNPs were then assigned chromosome positions and a sliding window method (window size of 50 SNPs; step size of one SNP) was adopted to identify recombination events for each individual. All the recombination sites were merged and sorted with 20 kb intervals^21^. In the end, the filtered 536,824 SNPs were combined into 2,013 bins. These were used to construct 11 linkage maps, resulting in 2180.14 cM spanning the whole genome. In addition, 1,556 scaffolds with 597.52 Mb were anchored, accounting for 94.42% of the assembled genome (Table 3).

### Transposable elements annotation

Transposable elements (TEs) annotation were performed by a combination of homology-based and *de novo* prediction approaches. Homology-based approach involved searching commonly used databases for known TEs at both DNA and protein level. With default parameters, RepeatMasker 3.3.0^22^ was used to identify TEs against the Repbase TE library 18.07^23^ and RepeatProteinMask^22^ was used to identify TEs at the protein level in the genome assembly. For *de novo* prediction, RepeatModeler software (http://www.repeatmasker.org/) was used in constructing the *de novo* repeat library. Tandem repeats were then predicted by TRF^24^ with parameters set to “Match = 2, Mismatch = 7, Delta = 7, PM = 80, PI = 10, Minscore = 50 and MaxPeriod = 2000’’. In total, we identified 294.95 Mb of the transposable elements, accounting for 46.47% of the asparagus bean genome (Table 4 and Table 5). Among all TEs, long terminal repeat (LTR), which are important determinants of angiosperm genome size variation, constituted 19.24% of the assembled genome. DNA TEs accounted for 7.2% of the total sequence.

### Gene annotation

We used *de novo*, homology and RNA-Seq-based prediction methods to annotate protein-coding genes in the asparagus bean genome. Three *de novo* prediction programs, Augustus^25^, Genscan^26^ and GlimmerHMM^26^ were used to annotate protein-coding genes while gene model parameters were trained from *Arabidopsis thaliana*. For homology-based prediction, protein sequences of all the protein-coding genes of eleven species including common bean (*Phaseolus vulgaris*), soybean)*Glycine max*), pigeonpea (*Cajanus cajan*), chickpea (*Cicer arietinum*), mungbean (*Vigna radiate*), adzuki bean (*Vigna angularis*), lotus (*Lotus japonicus*), medick (*Medicago truncatula*), Arabidopsis (*Arabidopsis thaliana*), grape (*Vitis vinifera*), and rice (*Orzya sativa*), were first mapped to the asparagus bean genome using TblastN with the parameter E-value=1e-05. GeneWise^27^ was then used to predict gene structure within each protein-coding region. RNA-Seq data of root and stem tissues^12^ were aligned to the asparagus bean genome using TopHat on default settings. Finally, the predicted genes were merged by EvidenceModeler (EVM)^28^ to generate a consensus and non-redundant gene set. This process produced 42,609 protein-coding genes with an average length of 3,156 bp (Table 6).

With BLASTP (E-value≤10 ^-5^), gene functions were assigned according to the best hit of alignment to SwissProt^29^, TrEMBL^30^, and KEGG^31^ database. Functional domains and motifs of asparagus bean genes were determined by InterProScan^32^, which analyzed peptide sequences against protein databases including SMART, ProDom, Pfam, PRINTS, PROSITE and PANTHER. Gene Ontology (GO) terms for each gene were extracted from the corresponding InterPro entries. The result showed that 75.40% (32,126) of the total genes were supported by TrEMBL, 56.22% (23,953) by Swiss-Prot, and 59.27% (25,254) by InterPro. In addition, 10,096 (23.69%) genes could not be functionally annotated with current databases (Table 7).

**Table 7.** Functional annotation of predicted genes in asparagus bean genome.

|  | Number | Percent (%) |
| --- | --- | --- |
| Total | 42,609 | ----- |
| InterPro | 25,254 | 59.27 |
| GO | 19,254 | 45.19 |
| KEGG | 18,372 | 43.12 |
| Swiss-Prot | 23,953 | 56.22 |
| TrEMBL | 32,126 | 75.4 |
| NR | 32,356 | 75.94 |
| Annotated | 32,513 | 76.31 |

The tRNA genes were identified by tRNAscan-SE software^33^ with default parameters. The rRNA genes were identified based on homology search to previously published plant rRNA sequences using BLASTN with parameters of “E-value=1e^-5^”. The snRNA and miRNA genes were identified by INFERNAL v1.0^34^ software against the Rfam database with default parameters. In all, 3,579 non-protein-coding genes were identified in the asparagus bean genome, including 1593 tRNAs, 1,076 rRNAs, 350 snRNAs, and 210 microRNAs (Table 8).

**Table 8.** Annotation of non-coding RNA in asparagus bean genome.

| Type |  | Copy(w) | Average Length(bp) | Total Length(bp) | % of genome |
| --- | --- | --- | --- | --- | --- |
| miRNA |  | 210 | 118.0571 | 24792 | 0.003906 |
| tRNA |  | 1593 | 75.10295 | 119639 | 0.018849 |
|  | rRNA | 538 | 155.6636 | 83747 | 0.013194 |
|  | 18S | 114 | 346.0877 | 39454 | 0.006216 |
| rRNA | 28S | 77 | 116.2338 | 8950 | 0.00141 |
|  | 5.8S | 22 | 146.8636 | 3231 | 0.000509 |
|  | 5S | 325 | 98.80615 | 32112 | 0.005059 |
|  | snRNA | 350 | 120.44 | 42154 | 0.006641 |
| snRNA | CD-box | 179 | 102.1285 | 18281 | 0.00288 |
|  | HACA-box | 24 | 123.7083 | 2969 | 0.000468 |
|  | splicing | 147 | 142.2041 | 20904 | 0.003293 |

### Code Availability

All tools used in this study were properly cited in the sections above. Settings and parameters were also clearly described.

## Data Records

The authors declare that all data reported here are fully and freely available from the date of publication. The Whole Genome Shotgun project has been deposited at GenBank (Data Citation 1). Raw read files are available at NCBI Sequence Read Archive (Data Citation 2 and Data Citation 3).

## Technical Validation

### DNA sample quality

DNA was quantified using 0.8% agarose gel electrophoresis and Qubit Fluorometer (Invitrogen, US). DNA concentrations were normalized to 100ng/µl for subsequent library construction.

### Assessment of the genome assembly and annotation

Completeness of the genome assembly was assessed with default settings using the Benchmarking Universal Single-Copy Orthologs (BUSCO)^35^ approach with a total of 1440 orthologue groups from the Embryophyta Dataset. The results showed that 93.2% of the core orthologs could be found in the asparagus bean genome, indicating a high-integrity assembly superior to the other four legume genomes. Furthermore, a previously reported high-density linkage map^5^ was used to assess the quality of anchored scaffolds. The sequences of 7,964 SNPs markers were aligned onto the 11 pseudo chromosomes using BLAT with parameters of “-fine”^36^. High accordance was shown between the assembled genome and the linkage map. We also aligned the raw reads from short insert-size sequencing back to the assembly and showed that approximately 94.88% of short reads could be successfully mapped.

### Comparison of asparagus bean genome with published common cowpea genomes

A comparison was performed (Table 9) between the asparagus bean genome and previously published common cowpea assemblies^7,9^. The asparagus bean genome assembly (549.81 Mb, non-N) was significantly larger than the first published IT97K-499-35^a^ genome^7^. Its size was close to the other two common cowpea survey assemblies (IT97K-499-35^b^ and IT86D-1010)^9^. The scaffold N50 size of our asparagus bean genome was the longest of all, reaching 2.7 Mb. Moreover, the asparagus bean assembly had about 94% of the scaffolds anchored into 11 pseudo-chromosomes according to the high-density genetic map. In addition, a set of 42,287 common cowpea coding sequences (CDS) derived from the single molecule real-time technology (*Vigna unguiculata* v1.0, http://phytozome.jgi.doe.gov/) could be blasted back to our asparagus bean genome with 90% similarity. All these results showed that the asparagus bean genome was of high quality.

**Table 9.**
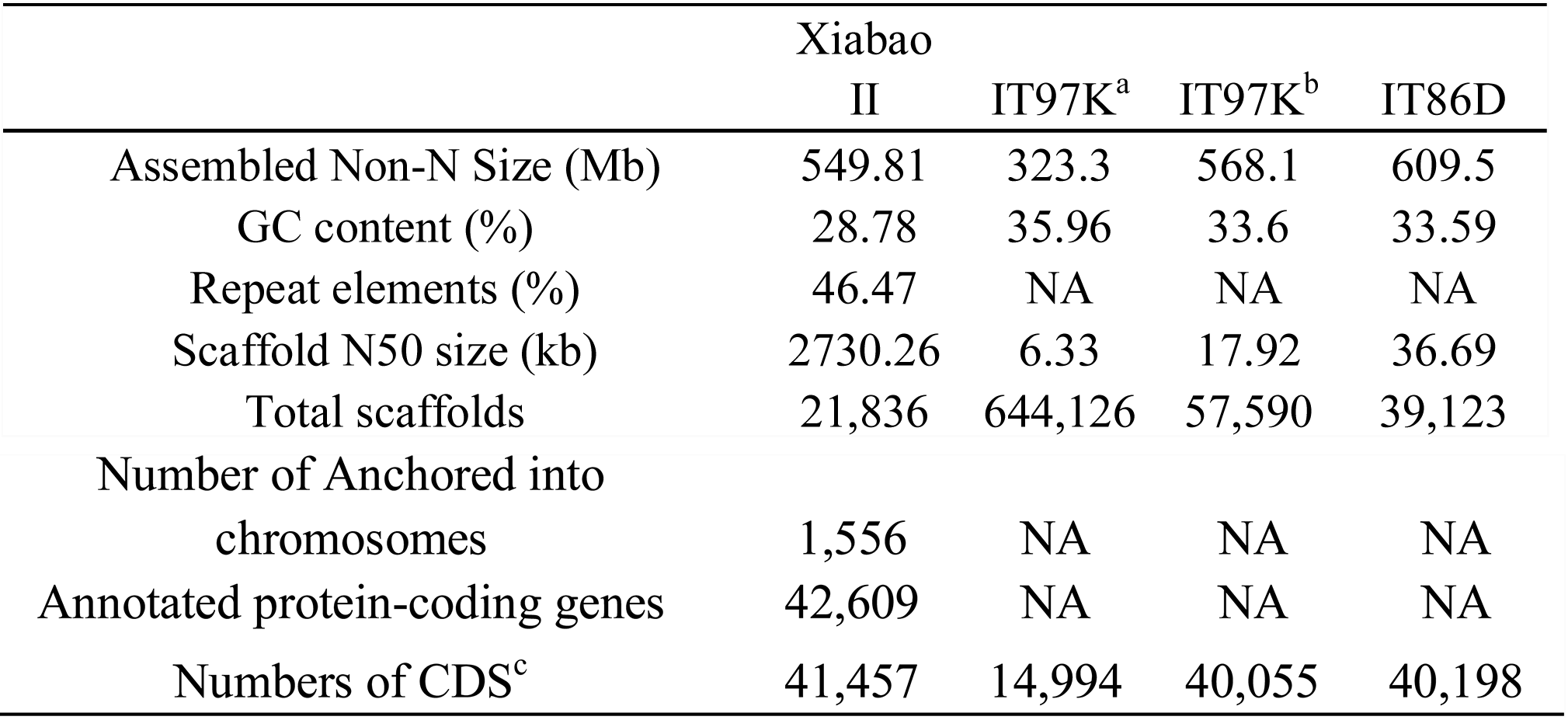
Comparisons of asparagus bean genome with published common cowpea assemblies.

## Acknowledgements

This study was funded by State Key Laboratory of Agricultural Genomics, BGI-Shenzhen, Shenzhen, China (NO.2011DQ782025). Financial support for *de novo* assembly was from the Key Lab of Agricultural Genomics, Chinese Ministry of Agriculture, BGI-Shenzhen, Shenzhen. China, Guangdong Provincial Key Laboratory of core collection of crop genetic resources research and application (NO.2011A091000047). Genetic map construction was supported by the fund from Shenzhen Municipal Government of China: Shenzhen Engineering laboratory of Crop Molecular design breeding. RNA-seq was supported by the fund from National Natural Science Foundation of China (NO.31501369).

## Author Contributions

Y.D., W.C, J.H.M., H.M.Y. and W.W. conceived and led the project. Y.D., X.M.N. and Y.L. contributed to secure funding. L.P. and C.Y.C. provided the sequencing samples and RNA-seq data. R.Z., Y.Z.W. and L.K. performed the sequencing. Q.J.X., R.Z. and Y.G. performed genome assembly and annotation. Q.J.X., X.D. and Z.Z. constructed the genetic map and anchored. Q.J.X., R.Z. and W.C. wrote the article.

## Competing Interests

The authors declare no competing interests.

## Data Citations

1. *GeneBank* PRJNA454850 (2018). *CNSA* CNP0000264 (2018).
2. *NCBI Sequence Read Archive* SRP144706 (SRR7135464-SRR7135488) (2018). *CNSA* CNP0000264 (2018).
3. *NCBI Sequence Read Archive* SRP144706 (SRR7125688-SRR7125784) (2018). *CNSA* CNP0000264 (2018).

## References

1. Ehlers, J.D., and Hall, A.E. Cowpea (Vigna unguiculata L. Walp.). Field Crops Research 53, 187–204.(1997).

2. Xu, P., Wu, X., Wang, B., Liu, Y., Qin, D., Ehlers, J.D., Close, T.J., Hu, T., Lu, Z., and Li, G. Development and polymorphism of Vigna unguiculata ssp. unguiculata microsatellite markers used for phylogenetic analysis in asparagus bean (Vigna unguiculata ssp. sesquipedialis (L.) Verdc.). Molecular Breeding 25, 675–684.(2010).

3. Fang, J., Chao, C.-C.T., Roberts, P.A., and Ehlers, J.D. Genetic diversity of cowpea [Vigna unguiculata (L.) Walp.] in four West African and USA breeding programs as determined by AFLP analysis. Genetic Resources and Crop Evolution 54, 1197–1209.(2006).

4. Jayathilake, C., Visvanathan, R., Deen, A., Bangamuwage, R., Jayawardana, B.C., Nammi, S., and Liyanage, R. Cowpea: an overview on its nutritional facts and health benefits. J Sci Food Agric 98, 4793–4806.(2018).

5. Xu, P., Wu, X., Muñoz-Amatriaín, M., Wang, B., Wu, X., Hu, Y., Huynh, B.L., Close, T.J., Roberts, P.A., Zhou, W., et al. Genomic regions, cellular components and gene regulatory basis underlying pod length variations in cowpea (V. unguiculata L. Walp). Plant Biotechnology Journal 15, 547–557.(2017).

6. Chen, X., Laudeman, T.W., Rushton, P.J., Spraggins, T.A., and Timko, M.P. CGKB: an annotation knowledge base for cowpea (Vigna unguiculata L.) methylation filtered genomic genespace sequences. BMC Bioinformatics 8, 129.(2007).

7. Munoz-Amatriain, M., Mirebrahim, H., Xu, P., Wanamaker, S.I., Luo, M., Alhakami, H., Alpert, M., Atokple, I., Batieno, B.J., Boukar, O., et al. Genome resources for climate-resilient cowpea, an essential crop for food security. Plant J 89, 1042–1054.(2017).

8. Muchero, W., Diop, N. N., Bhat, P. R., Fenton, R. D., Wanamaker, S., Pottorff, and M. e.a. A consensus genetic map of cowpea [Vigna unguiculata (L) Walp.] and synteny based on EST-derived SNPs. Proc. Natl. Acad. Sci. U.S.A., 18159–18164.(2009).

9. Spriggs, A., Henderson, S.T., Hand, M.L., Johnson, S.D., Taylor, J.M., and Koltunow, A. Assembled genomic and tissue-specific transcriptomic data resources for two genetically distinct lines of Cowpea (Vigna unguiculata (L.) Walp). Gates Open Research 2.(2018).

10. Xu, P., Wu, X., Wang, B., Liu, Y., Ehlers, J.D., Close, T.J., Roberts, P.A., Diop, N.N., Qin, D., Hu, T., et al. A SNP and SSR based genetic map of asparagus bean (Vigna. unguiculata ssp. sesquipedialis) and comparison with the broader species. PLoS One 6, e15952.(2011).

11. Kongjaimun, A., Kaga, A., Tomooka, N., Somta, P., Shimizu, T., Shu, Y., Isemura, T., Vaughan, D.A., and Srinives, P. An SSR-based linkage map of yardlong bean (Vigna unguiculata (L.) Walp. subsp. unguiculata Sesquipedalis Group) and QTL analysis of pod length. Genome 55, 81–92.(2012).

12. Pan, L., Wang, N., Wu, Z., Guo, R., Yu, X., Zheng, Y., Xia, Q., Gui, S., and Chen, C. A High Density Genetic Map Derived from RAD Sequencing and Its Application in QTL Analysis of Yield-Related Traits in Vigna unguiculata. Front Plant Sci 8, 1544.(2017).

13. Luo, R., Liu, B., Xie, Y., Li, Z., Lam, T.-W., and Wang, J. SOAPdenovo2: an empirically improved memory-efficient short-read de novo assembler. GigaScience 1:18.(2012).

14. Porebski, S., Bailey, L.G., and Baum, B.R. Modification of a CTAB DNA Extraction Protocol for Plants Containing High Polysaccharide and Polyphenol Components. Plant Molecular Biology Reporter 15 (1) :8–15.(1997).

15. Patel, R.K., and Jain, M. NGS QC Toolkit: A Toolkit for Quality Control of Next Generation Sequencing Data. PLoS ONE 7(2): e30619.(2012).

16. Boetzer, M., and Pirovano, W. SSPACE-LongRead: scaffolding bacterial draft genomes using long read sequence information. BMC Bioinformatics 15, 211.(2014).

17. Li, H., and Durbin, R. Fast and accurate short read alignment with Burrows-Wheeler transform. Bioinformatics 25, 1754–1760.(2009).

18. Li, H., Handsaker, B., Wysoker, A., Fennell, T., Ruan, J., Homer, N., Marth, G., Abecasis, G., Durbin, R., and Genome Project Data Processing, S. The Sequence Alignment/Map format and SAMtools. Bioinformatics 25, 2078–2079.(2009).

19. Behrend, A., Borchert, T., Spiller, M., and Hohe, A. AFLP-based genetic mapping of the“bud-flowering” trait in heather (Calluna vulgaris). BMC Genetics 14:64.(2013).

20. Wu, Y., Bhat, P.R., Close, T.J., and Lonardi, S. Efficient and accurate construction of genetic linkage maps from the minimum spanning tree of a graph. PLoS Genet 4, e1000212.(2008).

21. Ling, H.Q., Ma, B., Shi, X., Liu, H., Dong, L., Sun, H., Cao, Y., Gao, Q., Zheng, S., Li, Y., et al. Genome sequence of the progenitor of wheat A subgenome Triticum urartu. Nature 557, 424–428.(2018).

22. Tarailo-Graovac, M., and Chen, N. Using RepeatMasker to Identify Repetitive Elements in Genomic Sequences.(2009).

23. Bao, W., Kojima, K.K., and Kohany, O. Repbase Update, a database of repetitive elements in eukaryotic genomes. Mob DNA 6, 11.(2015).

24. Benson, G. Tandem repeats finder: a program to analyze DNA sequences. Nucleic Acids Research 2:573–580.(1999).

25. Stanke, M., Steinkamp, R., Waack, S., and Morgenstern, B. AUGUSTUS: a web server for gene finding in eukaryotes. Nucleic Acids Research 32, W309–W312.(2004).

26. Burge, C., and Karlin, S. Prediction of Complete Gene Structures in Human Genomic DNA. J. Mol. Biol 268, 78–94.(1997).

27. Birney, E., and Durbin, R. Using GeneWise in the Drosophila Annotation Experiment. Genome Research 10:547–548.(2000).

28. Haas, B.J., Salzberg, S.L., Zhu, W., Pertea, M., Allen, J.E., Orvis, J., White, O., Buell, C.R., and Wortman, J.R. Automated eukaryotic gene structure annotation using EVidenceModeler and the Program to Assemble Spliced Alignments. Genome Biol 9, R7.(2008).

29. Bairoch, A., and Apweiler, R. The SWISS-PROT protein sequence database and its supplement TrEMBL in 2000. Nucleic Acids Research 1:45–48.(2000).

30. Boeckmann, B. The SWISS-PROT protein knowledgebase and its supplement TrEMBL in 2003. Nucleic Acids Research 31, 365–370.(2003).

31. Kanehisa, M., and Goto, S. KEGG: Kyoto Encyclopedia of Genes and Genomes. Nucleic Acids Research 1:27-30.(2000).

32. Jones, P., Binns, D., Chang, H.Y., Fraser, M., Li, W., McAnulla, C., McWilliam, H., Maslen, J., Mitchell, A., Nuka, G., et al. InterProScan 5: genome-scale protein function classification. Bioinformatics 30, 1236–1240.(2014).

33. Schattner, P., Brooks, A.N., and Lowe, T.M. The tRNAscan-SE, snoscan and snoGPS web servers for the detection of tRNAs and snoRNAs. Nucleic Acids Res 33, W686–689.(2005).

34. Nawrocki, E.P., Kolbe, D.L., and Eddy, S.R. Infernal 1.0: inference of RNA alignments. Bioinformatics 25, 1335–1337.(2009).

35. Simão, F.A., Waterhouse, R.M., Ioannidis, P., Kriventseva, E.V., and Zdobnov, E.M. BUSCO: Assessing genome assembly and annotation completeness with single-copy orthologs. Bioinformatics 31, 3210–3212.(2015).

36. Kent, W.J. BLAT-the BLAST-like alignment tool. Genome Res 12, 656–664.(2002).

